# Murine leukemia virus can infect non-dividing cells

**DOI:** 10.1101/2023.08.30.555458

**Authors:** Karen Salas-Briceno, Wenming Zhao, Susan R. Ross

## Abstract

Retroviral reverse transcription starts within the capsid and uncoating and reverse transcription are mutually dependent. There is still debate regarding the timing and cellular location of HIV’s uncoating and reverse transcription and whether it occurs solely in the cytoplasm, nucleus or both. HIV can infect non-dividing cells because there is active transport of the preintegration complex (PIC) across the nuclear membrane, but Murine Leukemia Virus (MLV) is thought to depend on cell division for replication and whether MLV uncoating and reverse transcription is solely cytoplasmic has not been studied. Here, we used NIH3T3 and primary mouse dendritic cells to determine where the different stages of reverse transcription occur and whether cell division is needed for nuclear entry. Our data strongly suggest that in both NIH3T3 cells and dendritic cells (DCs), the initial step of reverse transcription occurs in the cytoplasm. However, we detected MLV RNA/DNA hybrid intermediates in the nucleus of dividing NIH3T3 cells and non-dividing DCs, suggesting that reverse transcription can continue after nuclear entry. We also found that the MLV PIC requires cell division to enter the nucleus of NIH3T3 cells. In contrast, we show that MLV can infect non-dividing primary DCs, although integration of MLV DNA in DCs still required the viral p12 protein. Knockdown of several nuclear pore proteins dramatically reduced the appearance of integrated MLV DNA in DCs but not NIH3T3 cells. Additionally, MLV capsid associates with the nuclear pore proteins NUP358 and NUP62 during infection. These findings suggest that simple retroviruses, like HIV, gain nuclear entry by traversing the nuclear pore complex in non-mitotic cells.

**Author Summary:** It is widely believed that gammaretroviruses like MLV require cell division to achieve nuclear entry and complete their replication. We show here that while this is true for rapidly dividing tissue culture cells, in quiescent cells like dendritic cells, the natural targets of MLV infection, the virus establishes infection without cell division. These studies show that the requirements for retrovirus infection depend on the cell type.

## Introduction

Retroviruses enter cells when the viral and host membranes fuse. Retroviruses replicate by reverse transcribing viral RNA into DNA using the virus-encoded reverse transcriptase (RT) [1]. Reverse transcription initiates within the capsid and capsid dissociation and reverse transcription are mutually dependent; a rigid capsid prevents reverse transcription from proceeding and the generation of DNA, which is more rigid than RNA, promotes capsid dissociation [2]. After reverse transcription and nuclear entry of the preintegration complex (PIC), which contains reverse-transcribed viral DNA in complex with host and viral proteins, the double-stranded DNA provirus integrates into the host genome via interaction of the viral integrase (IN) and host proteins like lens epithelium-derived growth factor (LEDGF; HIV) and bromodomain and extra terminal family (BET; MLV) [3-5].

Lentiviruses enter the nucleus of quiescent cells through interaction of viral capsid/nucleocapsid proteins with the nuclear pore complex (NPC) [6]. Retroviruses like murine leukemia viruses (MLV) and mouse mammary tumor virus (MMTV) require cell division and nuclear membrane breakdown for PIC entry because they are thought to lack viral proteins that interact with the NPC [6-10]. MLV, like HIV, dissociates its capsid to achieve integration [6]. The MLV p12 protein, encoded by the *gag* gene, tethers the PIC to chromosomes, allowing it to access chromosomes during mitosis [11]. Both MLV and MMTV infect non-dividing dendritic cells (DCs) *in vivo* [12-14]; how nuclear entry occurs in these cells is not known.

Recently, there has been debate as to where in the cell HIV reverse transcription occurs. Using HIV pseudoviruses bearing VSV G protein and tissue culture cells, several groups have shown that CA-containing reverse transcription complexes (RTC) enter the nucleus and that viral DNA synthesis is nuclear [15-18]. Capsid clearly contributes to HIV reverse transcription and integration; it has been demonstrated in a cell-free system that small patches of HIV-1 capsid loss are needed to accommodate the growing RTC and initiate integration [19]. Unlike the bullet shape of the HIV capsid, which is thought to facilitate nuclear pore entry, little is known about where reverse transcription occurs for other retroviruses including MLV, whose capsids are spheroid or polyhedral [9, 17].

Another unresolved issue for all retroviruses is the timing and location of host restriction during infection, particularly in natural targets of infection (e.g., lymphocytes, macrophages, dendritic cells) and how these factors coordinate with reverse transcription. Host anti-viral factors like apolipoprotein B mRNA editing enzyme catalytic subunit 3 (APOBEC3) proteins, which are packaged into retroviral virions, deaminate cytidine residues in the single-stranded DNA (ssDNA) generated by reverse transcription, thereby resulting in uracilated DNA, but whether this occurs in the nucleus, cytoplasm or both is not known. We recently showed that the nuclear base excision repair enzyme that removes uracil from ssDNA, uracil DNA glycosylase (UNG) works in the nucleus but not the cytoplasm to remove U residues from unintegrated MLV DNA deaminated by APOBEC3G, suggesting that at least some reverse transcription occurs in this compartment [20]. However, many sensors that detect HIV and MLV reverse transcripts, including cGAS, TREX1 and STING, are largely cytoplasmic and we and others have shown that sensing by these factors is sensitive to capsid stability [21-25]. Indeed, MLV encodes a protein called glycosylated Gag (glycoGag) protein that enhances capsid stability and prevents APOBEC3 and nucleic acid sensors from accessing the RTC [21, 22]. Similar results have been demonstrated for HIV - the HIV-2 capsid is more labile than the HIV-1 capsid and is more susceptible to nucleic acid sensors [23, 26].

Here we show that MLV requires cell division to efficiently enter the nucleus of NIH3T3 cells, although low levels of infection can be detected in quiescent cells. We also found that MLV RNA/DNA hybrid intermediates could be detected in the nucleus as well as the cytoplasm of dividing cells, suggesting that reverse transcription can occur after nuclear entry if reverse transcription is not complete prior to reformation of the nuclear envelope after cell division. In contrast to NIH3T3 cells, we found that MLV readily infected non-dividing primary dendritic cells and that early reverse transcripts could be equally detected in the nucleus and cytoplasm. Moreover, knockdown of several nucleoporins (NUPS) implicated in HIV trafficking across the nuclear envelope dramatically reduced the appearance of integrated MLV DNA in DCs but not NIH3T3 cells. These data suggest that MLV’s requirement for cell division is cell type-dependent and that in natural targets of infection, MLV PICs traffic across the nuclear membrane and efficiently establish infection.

## Results

### Incoming MLV capsid is found in the nucleus

Recent studies have demonstrated that HIV capsid can be detected in the nucleus at early times post-infection (reviewed in [27]). We first determined if MLV capsid (p30) could be found in the nucleus shortly after infection. NIH3T3 cells were infected with Moloney MLV and at 2- and 4-hours post-infection (hpi), the cells were fractionated and subjected to western blot analysis using anti-MLV p30 antibodies. As positive controls, NIH3T3 cells persistently infected with MLV were also tested. At 2 hpi, p30 was readily detected in the nuclear fractions and by 4 hpi, this level had decreased, likely coinciding with capsid dissociation (Fig. 1A). Nuclear p30 was also detected in persistently infected cells. Thus, like HIV, MLV capsid can be detected in the nucleus early after infection.

**Fig. 1.**
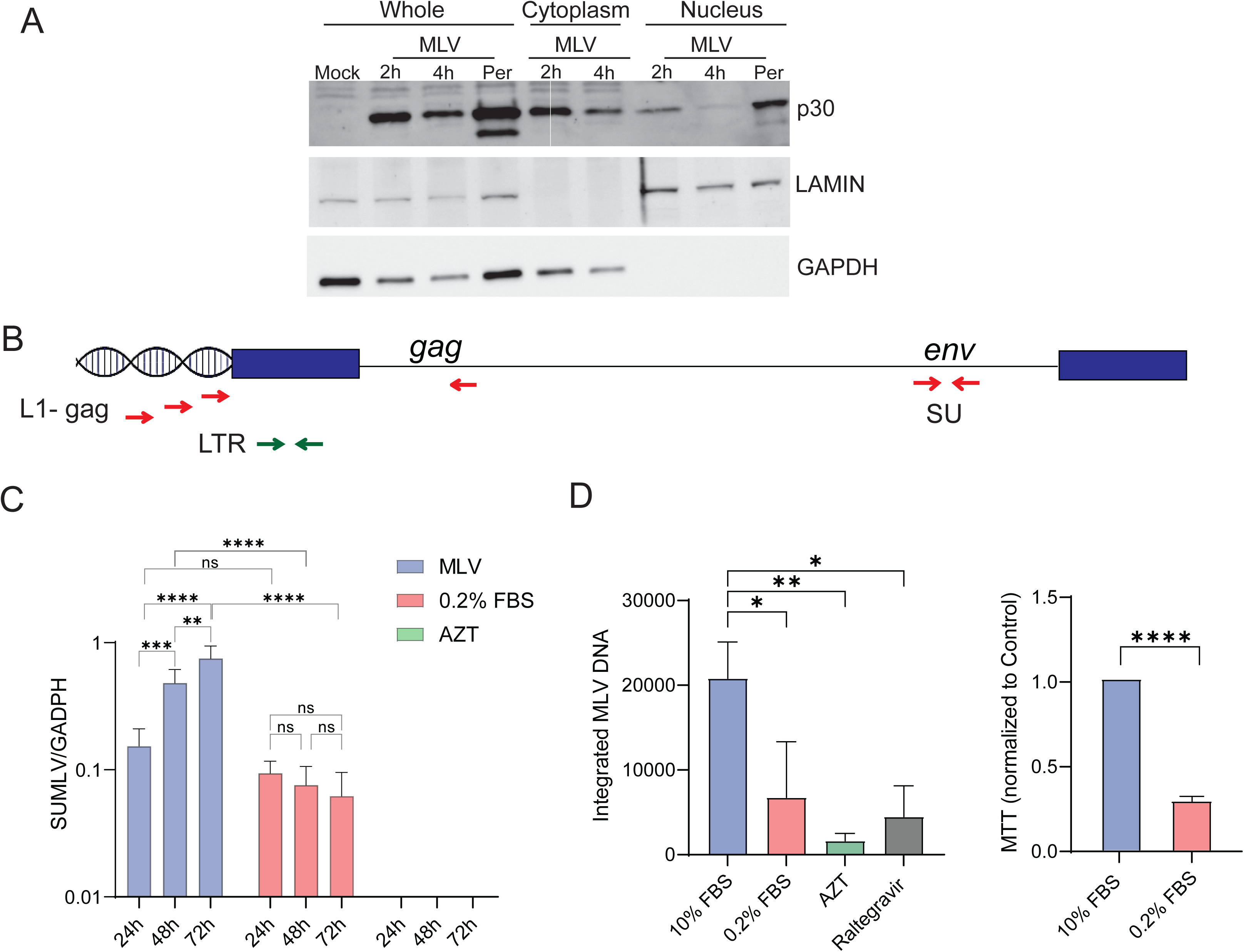
Efficient replication of MLV in NIH3T3 cells requires cells division. A) MLV (MOI = 1) was incubated on ice for 1 hr with NIH3T3 cells and warmed to 37°C for the indicated times. Western blots on fractionated extracts were performed with the indicated antibodies. NIH3T3 cells persistently infected with virus were used as positive controls. LAMIN B1 and GAPDH served as controls for nuclear and cytoplasmic fractions, respectively. This experiment was repeated 3 times with similar results. B) Map of the MLV genome and the primers used to detect integrated proviruses (round 1 PCR L1-gag/gag and round 2 nested PCR LTR primers) and virus infection levels (SU). C) NIH3T3 cells were grown in 0.2% or 10% serum, infected with MLV (MOI=1) and infection levels were measured at the indicated times. AZT treatment was used to block virus spread. D) Integrated MLV was determined in dividing, serum-starved and AZT- or raltegravir-treated cells. Shown to the right is the MTT assay, demonstrating that serum starvation decreased cell division. All experiments were performed 3 times. Shown is the average ± SD. Significance was determined by 2-way ANOVA, using Tukey’s multiple comparisons test (MLV in C and integrated MLV in D) or unpaired T test (MTT). \**P*≤ 0.03; \*\**P*≤ 0.002; *** *P*≤ 0.0003; \*\*\*\**P*≤ 0.0001.

### MLV reverse transcription intermediates are found in the nuclei of NIH3T3 cells

Many labs have studied MLV replication in growth-arrested, non-dividing tissue culture cells and shown that cell division is required for infection [6-8]. We also looked at virus spread in serum-starved and the reverse transcription inhibitor zidovudine/azidothymidine (AZT)-treated cells, by measuring the level of MLV envelope sequences in genomic DNA at different times post-infection (Fig. 1B). As has been shown previously, the level of MLV increased by 48 hpi in untreated cells, while in the serum-starved there was no spread at 48 and 72 hpi (Fig. 1C). AZT inhibited all virus spread.

We also determined that induction of quiescence by serum starvation blocked integration at 24 hpi. NIH3T3 cells were grown in 0.2% or 10% fetal calf sera for 48 hr and then infected with MLV. Integration was measured using a *Line-1*/*gag* nested PCR, similar to *Alu/gag* approach used to detect HIV integration (Fig. 1B) [20, 28]. Treatment of cells infected with MLV with AZT or the integration inhibitor raltegravir decreased integration by –13 and 5-fold, respectively. We found that serum starvation reduced but did not completely eliminate MLV integration (3-fold reduction; Fig. 1D). However, it is possible that the low-level integration detected in serum-starved cells resulted from incomplete inhibition of cell division, since the fold-reduction in cell division was ∼90% (Fig. 1D and S1 Fig). To try to improve the inhibition of cell division, we tried aphidicolin treatment to arrest cells, but also saw that a small percentage of cells continued to replicate (S1 Fig.). In addition, aphidicolin treatment was more toxic than serum starvation (S1 Fig.).

We next determined where reverse transcription intermediates were found in different compartments of dividing and quiescent cells. Untreated and serum starved NIH3T3 cells were infected with MLV and at 4 hpi, nuclear and cytoplasmic fractions were immunoprecipitated with S9.6 antibody, which recognizes RNA/DNA hybrids. The nucleic acid was eluted from the immunoprecipitates and subjected to reverse-transcribed PCR, using primers that recognize the first step of reverse transcription (P_tRNA_ and P_R_) and early reverse transcripts (strong-stop; P_R_ and P_U5_) (Fig. 2A). Western blots demonstrated that the nuclear and cytoplasmic fractionation was successful (Fig. 2B). In dividing cells, low amounts of the P_tRNA/_P_R_ product were detected in the nucleus, at about 10-fold lower levels than in the cytoplasm (Fig. 2C). However, strong-stop DNA was detected at almost equal amounts in nuclear and cytoplasmic fractions, suggesting that early reverse transcription is primarily cytoplasmic but that later steps can occur in the nucleus. In contrast, in serum-starved cells both the P_tRNA/_P_R_ and strong-stop DNA were greatly reduced in the nucleus.

**Fig. 2.**
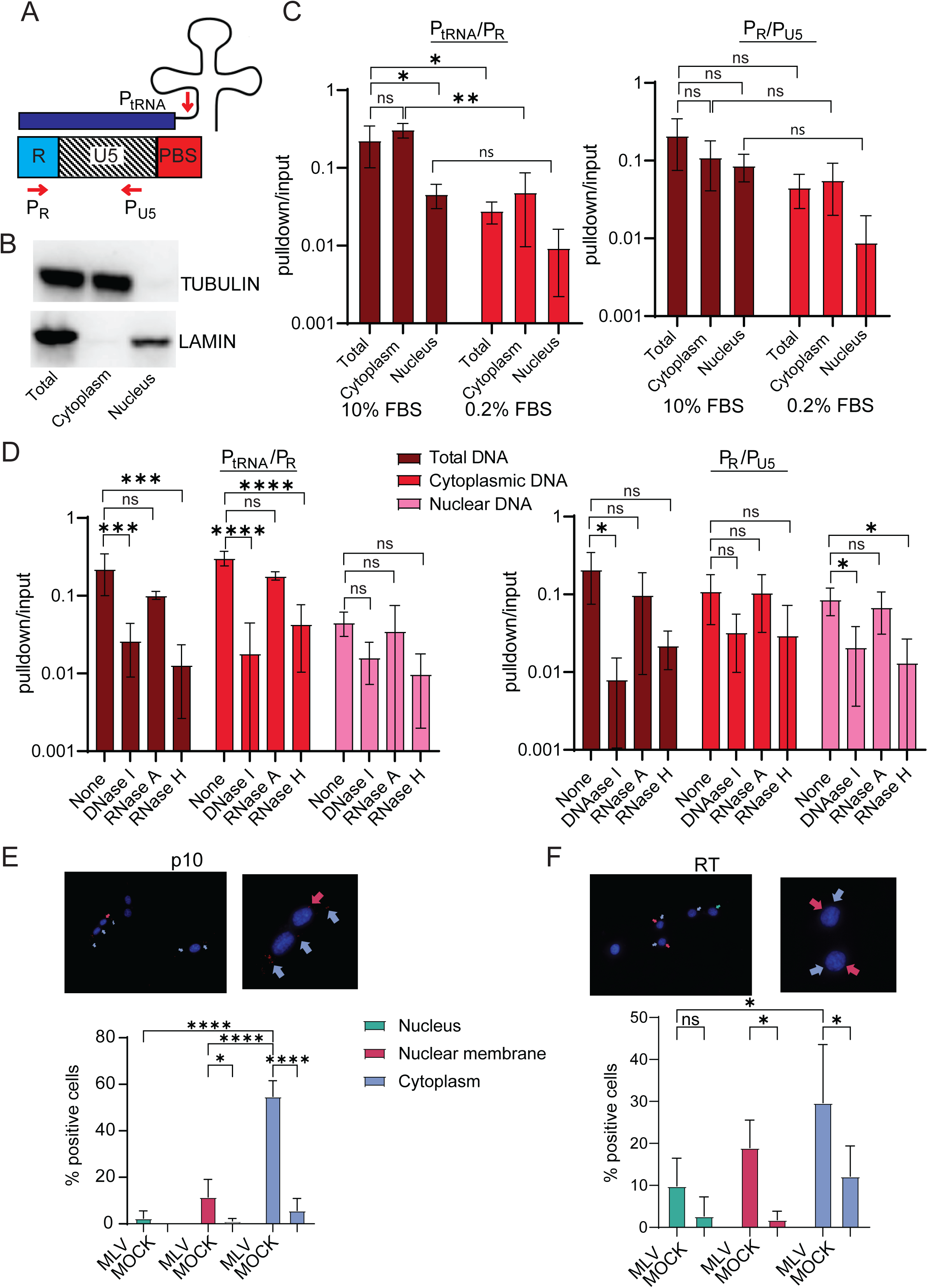
Early reverse transcripts (ERTs) are both nuclear and cytoplasmic. A) Map of the primers used to detect ERTs. B) Western blot showing cell fractionation. LAMIN B1 was used to identify nuclear and TUBULIN cytoplasmic fractions. C) Nuclear, cytoplasmic and total extracts from dividing and serum-starved NIH3T3 cells infected with MLV (MOI=1) were immunoprecipitated with antibody S6, the nucleic acid was released from the immunoprecipitates and subjected to qPCR with primers that detect the 1st step of reverse transcription (P_tRNA_/P_R_) and strong stop DNA (P_R_/P_U5_). Shown is the average of 3 experiments ± SD. D) The immunoprecipitates from (C) were treated with the indicated enzymes prior to qPCR. Shown is the average of 3 experiments ± SD. E) and F) PLA was performed using antibodies to MLV p10 (E) or RT (F) and S6. Shown below the photographs is quantification of the appearance of the PLA+ spots in different cellular locations (25 fields; 210 – 270 cells). Shown is the average number for each location per field ± SD. Significance was determined by 1-way ANOVA. \**P*≤ 0.04; \*\**P*≤ 0.005; \*\*\**P*≤ 0.0003; \*\*\*\**P*≤ 0.0001.

To ensure that the immunoprecipitated products were RNA/DNA hybrids, we treated the fractions with DNaseI, RNaseH and RNaseA. Only RNaseH and DNaseI abolished the PCR products from cells grown in 10% and 0.2% serum, confirming that the products were early reverse transcripts (Fig. 2D). Taken together, these data show that cell replication is needed for efficient trafficking of the PIC into the nucleus of NIH3T3 cells but suggest that partially reverse-transcribed replication products can complete replication in the nucleus after cell division finishes.

We also used proximity ligation assays (PLA) to examine the co-localization of reverse transcripts and MLV proteins, using the S9.6 antibody and anti-MLV antibodies. DNA/RNA hybrids co-localized with MLV nucleocapsid (NC; p10) at the nuclear membrane, and cytoplasm of infected but not uninfected cells; low levels of co-localization were also seen in the nucleus (Fig. 2E). Similar results were seen with the anti-RT antibodies, although there were also PLA positive signals in the cytoplasm of uninfected cells (Fig. 2F; S2 Fig.). Previous studies have suggested that NIH3T3 cells harbor endogenous MLVs, which may be the source of this signal [29, 30].

These data show that reverse transcription can occur in the nuclei of non-dividing NIH3T3 cells, but that it is inefficient and confirm previous findings that MLV replication in tissue culture cells requires cell division.

### Non-dividing DCs are highly infected with MLV

We previously showed that DCs can be infected with both MMTV and MLV *in vivo* and *in vitro* [12, 14]. Once differentiated, DCs are non-replicating. We next tested whether DCs isolated from bone marrow (BMDC) could be infected with MLV. Because we showed previously that mouse APOBEC3 can reduce early reverse transcription of incoming virus, we used BMDCs isolated from APOBEC3 knockout mice for these experiments to increase sensitivity; NIH3T3 cells do not express APOBEC3 [31-33]. We first confirmed that >98% of DCs analyzed at 10 – 11 days of differentiation ex vivo were quiescent (Fig. 3A). These cells were then infected with MLV and at 24 hpi, genomic DNA was analyzed for integrated MLV. Integrated DNA was easily detected, and the levels were reduced by treatment with AZT and raltegravir (Fig. 3B).

**Fig. 3.**
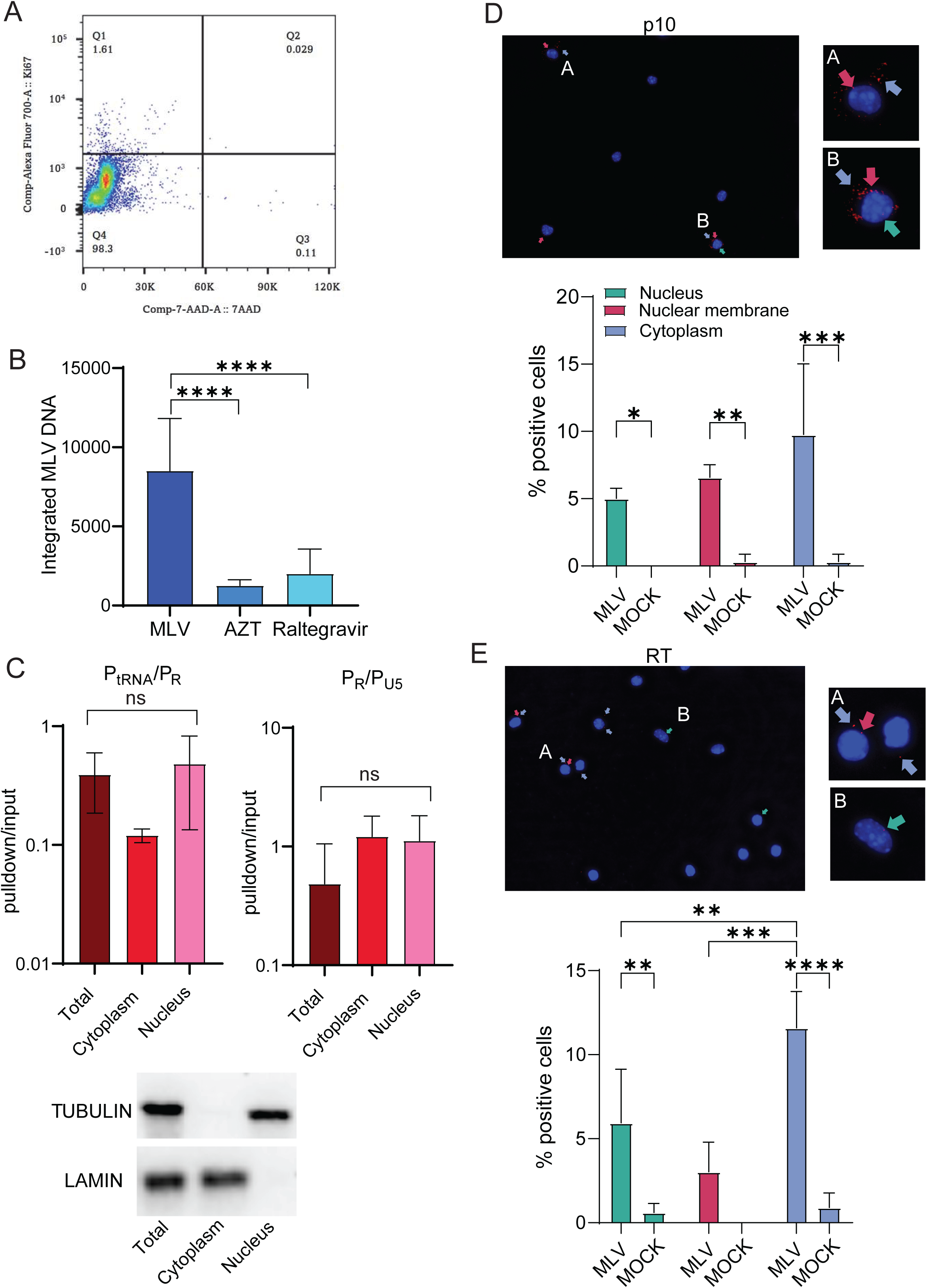
Primary DCs are readily infected with MLV. A) Representative FACS plot demonstrating that primary DCs are non-dividing. B) BMDCs were infected with MLV (MOI=1) and integrated proviral DNA was measured. AZT and raltegravir treatment were used as controls to inhibit infection. Shown is the average of 3 independent experiments ± SD. C) Nuclear, cytoplasmic and total extracts from BMDCs were immunoprecipitated with antibody S6 and subjected to qPCR with primer pairs P_tRNA_/P_R_ and P_R_/P_U5_. Experiments were performed a minimum of 3 times. Shown is the average ± SD. Significance was determined by 1-way ANOVA. ns, not significant. Shown below the graphs is a representative western blot showing the fractionation purity. LAMIN B1 was used to detect nuclear and TUBULIN to detect cytoplasmic fractions. D) and E) PLA was performed using antibodies to MLV p10 (D) or RT (E) and S6. Shown below the photographs is quantification of the appearance of the PLA+ spots in different cellular locations (25 fields; 480 – 570 cells). Shown is the average number for each condition per field ± SD. Significance was determined by 2-way ANOVA. \**P*≤ 0.04; \*\**P*≤ 0.002; \*\*\**P*≤ 0.0004; \*\*\*\**P*≤ 0.0001.

Next, to determine where early reverse transcription was occurring, we infected DCs and at 4 hpi, carried out fractionation followed by pulldown with S9.6 antibody and analysis of the bound nucleic acid. Unlike what we saw with NIH3T3 cells, approximately equal levels of both the P_tRNA/_P_R_ and P_tRNA/_P_U5_ RT-PCR products were found in the nucleus and cytoplasm (Fig. 3C). This suggested that MLV reverse transcription occurred in both compartments of infected DCs. We also found, using PLA, that RNA/DNA hybrids co-localized with MLV p10 at the nuclear membrane and the cytoplasm and nucleus of DCs at almost equal levels (Fig. 3D). Similar results were seen with co-localized RT (Fig. 3E). As we saw with NIH3T3 cells, a PLA signal with anti-RT and S9.6 antibodies could also be detected in uninfected cells.

### MLV p12 is required for BMDC infection

The MLV *gag* gene encodes a protein, p12, which is required for anchoring of viral dsDNA to the chromosomes, thereby facilitating integration in dividing cells [11, 34, 35]. To determine if p12 was also required for integration in non-dividing BMDCs, we infected them with mutant p12-M63-PM15, in which the C-terminal residues 70RREPP74 were replaced with alanines (70AAAAA74); this mutation abrogates chromatin binding of the PIC in dividing cells [35]. For comparison, we used pNCA, the cloned parental virus from which p12-M63-PM15 was constructed. No integrated DNA could be detected in p12-M63-PM15-infected cells at 24 hpi, while pNCA infected BMDCs to a similar extent as wild type MLV (Fig. 4A). The p12-M63-PM15 PIC did enter the nucleus, however, because 2-LTR circles, the nuclear by-product of unintegrated reverse-transcribed DNA, were detected at about 2-fold higher levels than with MLV or pNCA MLV, as expected for viral DNA that is unable to integrate (Fig. 4B). Moreover, RNA/DNA early reverse transcription products were equally detected in the cytoplasm and nucleus in MLV-, pNCA MLV- and p12-M63-PM15-infected BMDCs at 4 hpi (Fig. 4C). Thus, like dividing cells, MLV p12 is required for integration of viral DNA into the chromosomes.

**Fig. 4.**
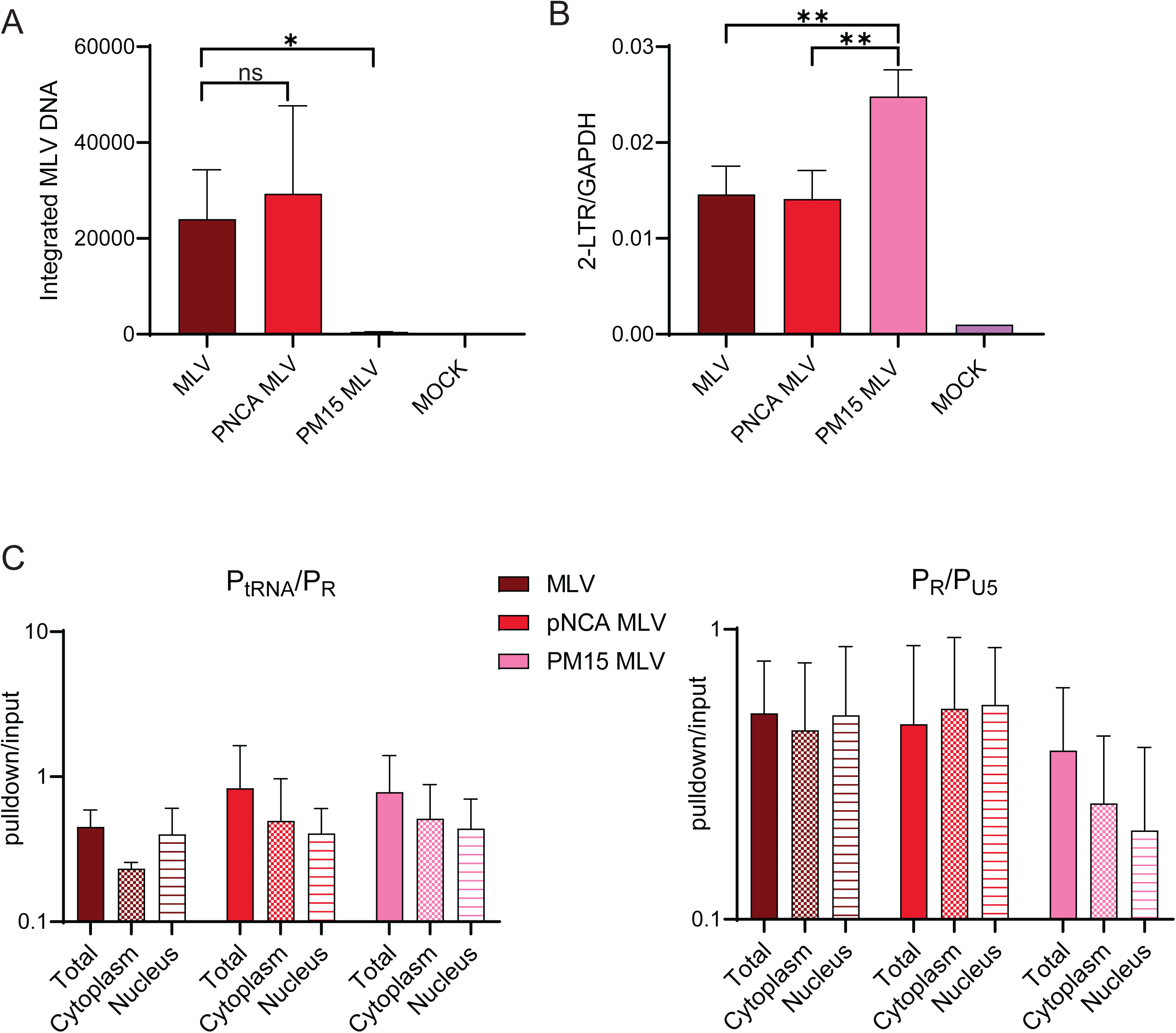
MLV p12 is needed for integration in BMDCs. A) Integrated MLV was determined in MLV wildtype (MLV and pNCA MLV)- and p12 mutant MLV (PM15)-infected cells. All experiments were performed 3 times. Shown is the average ± SD. Significance was determined by unpaired T test. \**P*≤ 0.0167. B) Nuclear DNA was amplified with primers that detect 2-LTR circles. Experiments were performed 3 times. Significance was determined by 1-way ANOVA. \*\**P*≤ 0.01. C) Nuclear, cytoplasmic and total extracts from BMDCs infected with the indicated virus were immunoprecipitated with antibody S6, the nucleic acid was released from the immunoprecipitates and subjected to qPCR with primers that detect the 1st step of reverse transcription (P_tRNA_/P_R_) and strong stop DNA (P_R_/P_U5_). Shown is the average of 3 experiments ± SD. No significant difference was seen between any of the groups using 1-way ANOVA.

### MLV infection of DCs relies on nucleoporins

It is well-established that the HIV PIC relies on the nuclear pore for entry into the nucleus of quiescent cells. Several nucleoporins have been implicated in this transport, including NUP358, NUP62, NUP88, NUP153 and NUP214 [36, 37]. To determine if any of these NUPs were required for MLV nuclear entry, we used siRNA knockdown in NIH3T3 cells (Fig. 5A) and BMDCs (Fig. 5B) to deplete their expression. As controls, AZT and raltegravir were used to block infection. In NIH3T3 cells, siRNA knockdown of these 5 nucleoporins had no effect on MLV infection at 24 hpi, as integrated provirus levels were equal to cells treated with the control siRNA (Fig. 5A). In contrast, knockdown of all 5 nucleoporins dramatically reduced infection in BMDCs, to similar levels seen with AZT and raltegravir treatment (Fig. 5B). Knockdown efficiency was similar in both cell types, and cell viability was not affected, as determined by MTT assays (Fig. 5A and 5B). By examining the fluorescence intensity in cells immunostained for each of the nucleoporins, we also found that knockdown of most reduced the levels of the others, with the exception of NUP358 and NUP88 (Fig. 5C and S3 Fig.). However, NUP358 and NUP88 knockdown was less efficient at the protein level than the other NUPS so this might account for the lack of effect on expression of other NUPs (Fig. 5C).

**Fig. 5.**
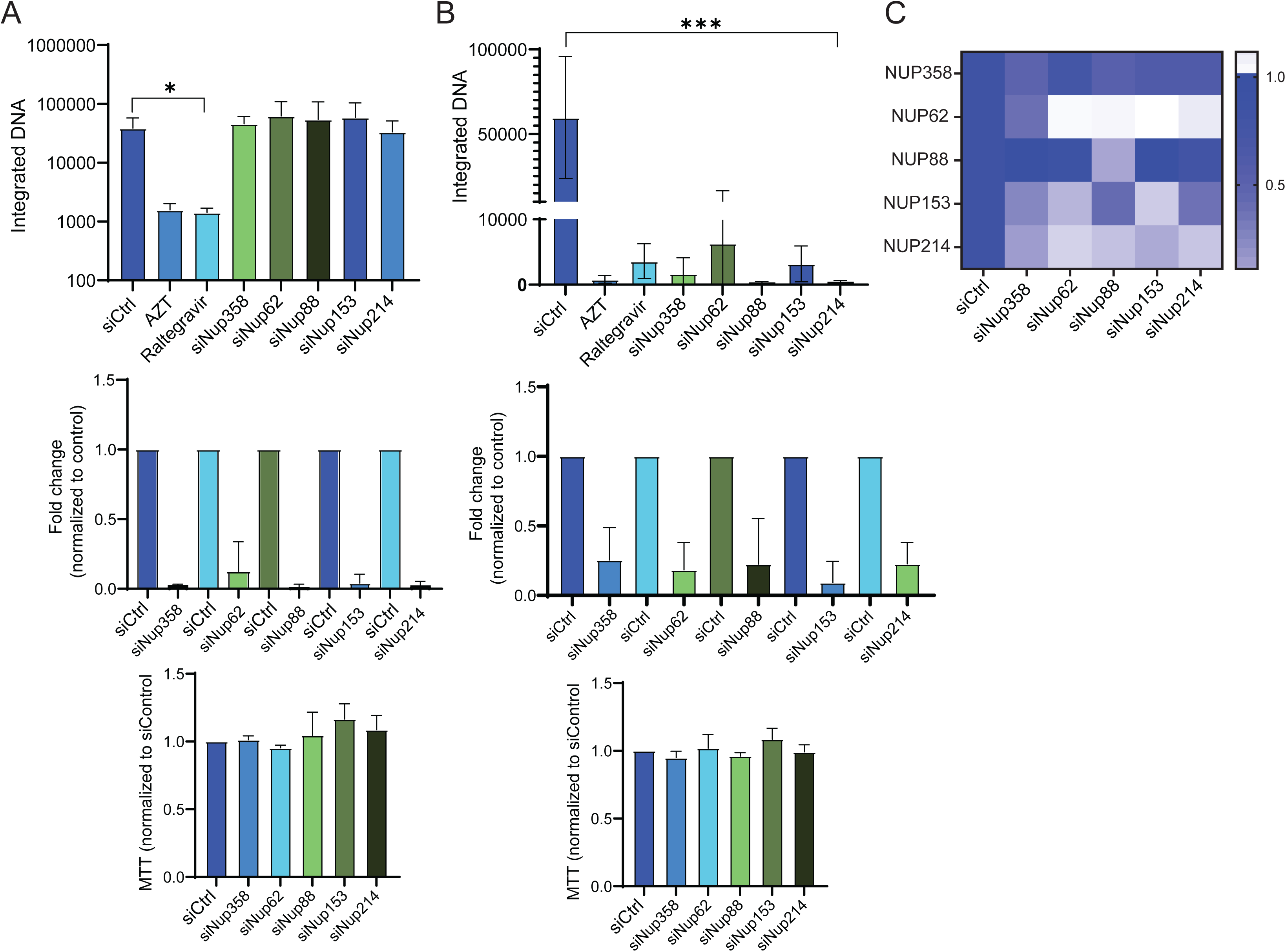
Depletion of NUP proteins decreases MLV infection of DCs but not NIH3T3 cells. A) siRNA-mediated knockdown of the indicated NUPs in NIH3T3 cells. AZT and raltegravir treatment serve as controls. Knockdown efficacy is shown in the middle panel and effects on cell viability in the bottom panel. B) The same experiment was performed in BMDCs. All experiments were performed a minimum of 3 times. Shown is the average ± SD. Significance was determined by 1-way ANOVA. \**P*≤ 0.03; \*\*\**P*≤ 0.0009. C) The intensity of immunostaining for the indicated proteins after NUP knockdown was quantified. One hundred to 160 cells were analyzed for each knockdown condition. Representative images are shown in S3 Fig.

We next used PLA to examine whether MLV capsid co-localized with any of the NUPs. We were able test 2 nucleoporins, NUP358, which is located on the cytoplasmic side of the nuclear pore and NUP62, found within the channel, with anti-capsid antibody. To first examine interaction, we used a NIH3T3 cell line stably transfected with a CMV-driven gag/pol construct [13]. Many cells showed multiple co-localization spots with both nucleoporins (Fig. 6A). The PLA spots detected with NUP358 and NUP62 and CA antibodies localized to the cytoplasm, nuclear membrane and nucleus (Fig. 6A).

**Fig. 6.**
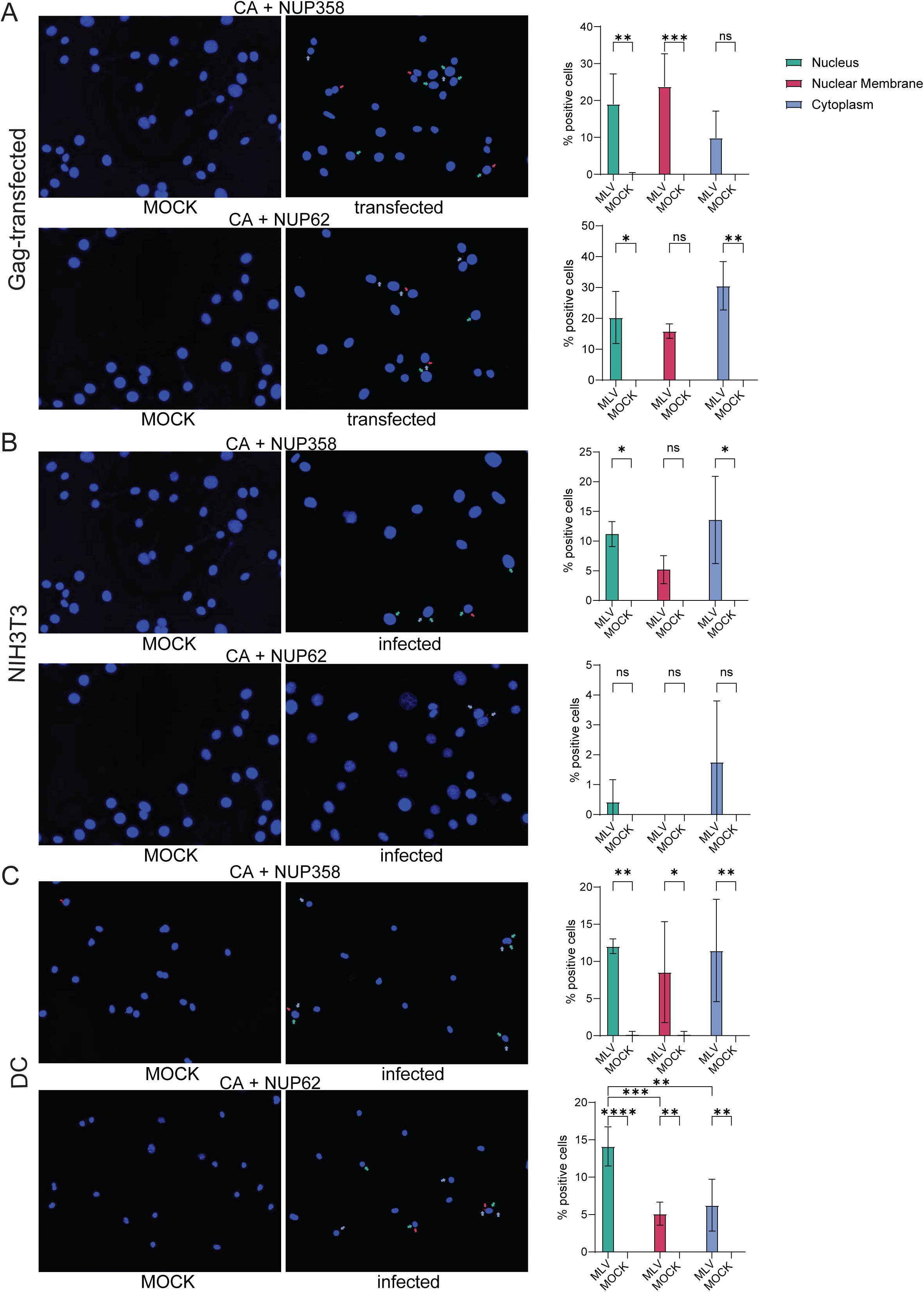
MLV CA co-localizes with NUP358 and NUP62. PLA was performed with anti-MLV CA and anti-NUP358 or -NUP62 antibodies. A) NIH3T3 cells transfected with a Gag expression plasmid. B) NIH3T3 cells at 4 hpi. C) BMDCs at 4 hpi. Shown to the right is the percentage of PLA spots per field for each condition (15 fields/condition; 115 – 320 cells; average ± SD). Significance was determined by 2-way ANOVA (Tukey’s multiple comparison test). \**P*≤ 0.05; \*\**P*≤0.01: \*\*\**P*≤0.001; \*\*\*\**P*≤ 0.0001.

We next examined NUP co-localization in MLV-infected NIH3T3 cells and BMDCs at 4 hpi. In both cases, co-localization of capsid was detected with both NUP358 and NUP62 (Fig. 6B and 6C). Interestingly, while NUP62/capsid was found inside the nucleus in infected BMDCs, in infected NIH3T3 cells the spots were mainly seen in the cytoplasm. This supports the preceding evidence that nuclear entry of MLV PICs largely requires cell division in NIH3T3 cells but not DCs.

## Discussion

Historically, it was thought that retroviral reverse transcription occurred entirely in the cytoplasm. However, recent studies have demonstrated HIV reverse transcription happens at least partially in the nucleus, although whether it initiates in this location is still under debate [15, 18, 27, 38, 39]. Here we showed that MLV reverse transcription occurs in the cytoplasm and nucleus of both dividing and non-dividing cells. In dividing cells, like NIH3T3 cells, we found that most reverse transcription is cytoplasmic, but that it can be completed in the nucleus. Since blocking NIH3T3 cell division greatly decreased the appearance of RTCs in the nucleus, nuclear reverse transcription likely occurs when the RTC ends up in the nucleus post-cell division. These results also explain our previous findings in 293T cells, suggesting that RNA/DNA intermediate products of reverse transcription were nuclear; we showed that although APOBEC3G largely deaminates cytoplasmic MLV reverse transcripts, the nuclear enzyme UNG, which preferentially removes uracils found in ssDNA, acts on unintegrated nuclear viral DNA [20].

Even in serum starved NIH3T3 cells, there were low levels of RTCs found in the nucleus and integration and infection was reduced by not completely eliminated. Because we were unable to complete arrest cell division, either by serum starvation or aphidicolin treatment, we cannot rule out that this low-level infection is due to a small percentage of dividing cells. We also found that serum starved NIH3T3 cells had lower overall levels of reverse transcription even in the cytoplasmic fraction, which may be due to the diminution of other host factors needed for MLV entry, uncoating or reverse transcription (Fig. 2B).

Many studies have shown that gammaretroviruses like MLV require cell division for viral replication, although these were performed in replication competent tissue culture cells and largely used virus spread as the readout for infection [6-10]. Here we show that in contrast to NIH3T3 cells, MLV PICs traffic across the nuclear membrane and efficiently infect non-dividing primary DCs. Whereas in NIH3T3 cells the appearance of RNA/DNA hybrids in the nucleus was greatly reduced, in DCs equal levels were seen in the nucleus and cytoplasm. Similarly, p10 in particular was found at higher levels in association with RNA/DNA hybrids in the nuclei of DCs compared to NIH3T3 cells. The RT/RTC association also was lower in the nuclei and cytoplasm of NIH3T3 cells compared to DCs, although the difference was less pronounced than with p10. This may be because the nucleocapsid is more dissociated from PICs found in NIH3T3 nuclei than in DCs.

It is known that the nuclear pore complex mediates nucleo-cytoplasmic transport of macromolecules and is required for the trafficking of viral PICs and as a result, replication of HIV in quiescent cells (reviewed in [27, 36]). In support of the trafficking of MLV RTCs/PICs in quiescent DCs compared to NIH3T3 cells, knockdown of various nucleoporins decreased MLV integration in the former but not the latter cell types. In addition, we were able to detect MLV capsid in the nucleus as early as 2 hpi, suggesting that some CA remains associated with the RTC following its translocation into the nucleus. Thus, like HIV, MLV capsid may be uncoated in two stages. The first stage is the loss of integrity of the intact core. This event seems to occur certainly in the cytoplasm but does not remove all the CA from the RTC prior to nuclear import [40].

Our results also indicate that MLV CA interacts with one or more of the same nucleoporins as HIV. In knockdown experiments, we were unable to directly assess which nucleoporin was critical, as knockdown of each nucleoporin altered the protein levels of the others. In the most dramatic case, *Nup62* knockdown decreased the expression of all other tested nucleoporins. We also observed that even though *NUP358* knockdown was not complete, it affected the MLV integration levels, indicating that even small disturbances in the nuclear pore alter the efficiency of MLV PIC nuclear entry.

Finally, using PLA assays we showed that in both NIH3T3 cells and DCs, MLV capsid interacts with NUP358 and NUP62 (Fig. 6C). NUP358 is found at the cytoplasmic side of the nuclear pore, while NUP62 is found in the channel [41]. We found that NUP358/CA interaction occurred equally in the cytoplasm and nucleus in both cell types, while NUP62/CA occurred in the nucleus and at the nuclear membrane in DCs but not NIH3T3s, where it was mostly seen in the cytoplasm. Although some MLV capsid and NUP358 and NUP62 interactions were seen in the cytoplasm, in most cases the co-localization was perinuclear. Whether this cytoplasmic interaction is the result of MLV causing NUP mislocalization is not clear. However, several other viruses, including HIV, have been shown to cause redistribution of NUPs to the cytoplasm (reviewed in [42]). Additional studies are also necessary to determine if MLV CA physically binds to multiple nucleoporins, and if MLV interaction with nucleoporins affects only nucleocytoplasmic trafficking, or also affects MLV’s access to the host chromatin and chromosomal site selection for integration. However, it is clear that even in nondividing DCs, the MLV p12 protein is still required for integration of viral DNA into the chromosomes.

Thus, our work shows that the gammaretrovirus MLV can infect non-dividing cells like DCs, thought to be the initial *in vivo* targets of MLV [12, 13, 43]. Interestingly, previous studies have suggested that gammaretroviruses can infect other quiescent cell types, such as neurons [44]. Our studies also suggest that reverse transcription and partial uncoating of MLV virions occur in the cytoplasm. This supports the observation that APOBEC3 deaminates cytoplasmic reverse transcripts and that cytoplasmic nucleic acid sensors respond to MLV RNA/DNA hybrids and reverse transcribed DNA [13, 20, 22, 45]. This action by anti-viral cytoplasmic proteins is important for host defense, particularly in sentinel cells of the immune system like DCs. Perhaps the relatively low immune response to retroviruses compared to other RNA viruses is because of the rapid movement of RTCs to the nucleus in sentinel cells, thereby avoiding cytoplasmic detection.

## Methods

### Cell cultures

NIH3T3 cells were cultured in Dulbecco’s Modified Eagle Medium (DMEM) supplemented with 10% FBS, 2mM L-glutamine (Gibco), 100 U/ml penicillin (Gibco), and 100 μg/ml streptomycin (Gibco). Cell cultures were maintained at 37°C with 5% CO2. For the BMDCs cultures, bone marrow from APOBEC3 knockout mice was harvested from the hind limbs of mice, cultured in RPMI supplemented with 10% FBS, 2 mM L-glutamine, 100 U/ml penicillin, 100 μg/ml streptomycin, and 50 μM β-mercaptoethanol and differentiated with 20 ng/ml murine granulocyte-macrophage colony-stimulating factor (GM-CSF) (Peprotech, 315-03). BMDCs were stained for CD11c and analyzed by flow cytometry (LSR-Fortessa flow cytometer; BD Biosciences). DC cell populations resulted in cultures that were >80% pure (S4 Fig.).

### Cell cycle analysis by flow cytometry

NIH3T3 cells and BMDCs were incubated in Fc block (BD Biosciences) in 2% FBS in PBS and fixed and permeabilized with BD Cytofix/Cytoperm™ Fixation/Permeabilization Kit (BD Biosciences). Then, cells were stained with 7-AAD Viability staining solution (BioLegend), Zombie Aqua Dye (BioLegend), and the antibody Alexa Fluor 700 anti-mouse Ki-67. Analysis was done in the LSR-Fortessa flow cytometer (BD Biosciences) using FlowJo v10 software (Tree Star, Inc.).

### Virus isolation

Moloney MLV was isolated from the supernatants of stably infected NIH 3T3 cells (cells in which infection is allowed to spread to 100% of the culture and maintained in this state thereafter). pNCA MLV and p12-M63-PM15 plasmids were a gift from Monica Roth [35]. The pNCA MLV and p12-M63-PM15 plasmids were transfected into 293T cells using Lipofectamine 3000 (Invitrogen). The media of the transfected cells were harvested 48 h post-transfection. Supernatants from NIH3T3 and 293T cells containing MLV were passed through a 0.45-mm filter, treated with 20 U/ml DNase I (Sigma) (NIH3T3 cells) or salt active nuclease (SERVA) (transfected 293T cells) at 37°C for 30 min, and centrifuged through a 25% sucrose cushion. After resuspension, titers of MLV and pNCA MLV were determined on NIH 3T3 cells. All viruses were subjected to reverse transcriptase quantitative PCR (RT-qPCR), and the number of viruses was estimated by standard curve analysis from the amount of virus-specific RNA, using primers located in the env gene (Table 1). Equal amounts of virus, normalized by RNA levels, were also analyzed by Western blots (S5 Fig.).

### Western blot analysis

Western blots analysis. For fractionation assays, NIH-3T3 cells were collected in ice-cold PBS, and fractionated into nuclear, cytoplasmic, and whole cell extracts by the Rapid, Efficient, and Practical (REAP) method (39). Cellular and viral roteins were detected by Western blots. Polyclonal goat anti-MLV antibody (NCI Repository) (2), anti-GAPDH (D16H11, Cell Signaling Technology/CST), anti-α-tubulin (T6074, Sigma-Aldrich), anti-lamin B1 (D4Q4Z, CST), horseradish peroxidase (HRP)-conjugated anti-rabbit (7074, CST), and (HRP)-conjugated anti-goat (A8919, Sigma-Aldrich) were used for detection, using Pierce ECL Western blotting substrate (Thermo Fisher Scientific, 32209).

### Real-time qPCR

RT-qPCRs were performed with MLV SU or 2-LTR primers (Table 1) using a Power SYBR green PCR kit (Promega) and the QuantStudio 5 real-time PCR system (Applied Biosystems), as previously reported [20]. DNA quantifications were normalized to glyceraldehyde-3-phosphate dehydrogenase (GAPDH). The amplification conditions were 50°C for 2 min, 95°C for 10 min, 40 cycles of 95°C for 15 s, and 60°C for 1 min. The efficiency of amplification was determined for each primer pair by generating a standard curve with 10-fold serial dilutions of a known concentration of DNA. For each primer pair, a no-template control was included, and each sample was run in triplicate. Levels of integrated MLV were determined by LINE1-gag nested qPCR, based on the Alu-gag qPCR for HIV. Briefly, 50 ng of total DNA was used to perform a PCR using a forward primer that targeted genomic LINE 1 sequences located randomly near integrated proviruses and an MLV-specific gag reverse primer (Table 1). The PCR product was diluted 10-fold, and 2.4 µl was used as input for the second qPCR, which was performed using MLV long terminal repeat (LTR) primers (Table 1) [20].

### Nucleic acid pulldowns

NIH3T3 and BMDC cells were infected with MLV (genome equivalent of a multiplicity of infection [MOI] of 1) in the presence of 8 mg/ml Polybrene (Sigma-Aldrich), and cells were incubated on ice for 1 h to allow virus binding. Cells were washed in cold phosphate-buffered saline and incubated at 37°C for 4 h. At 4 hpi the cells were fractionated by REAP method [46]. Then each fraction was incubated overnight with anti-DNA-RNA Hybrid, clone S9.6 Ab (Sigma-Aldrich), and G/A-agarose beads (Santa Cruz). The beads were washed with high-salt buffer (25mM Tris-HCl, pH 7.8, 500mM NaCl, 1mM EDTA, 0.1% SDS, 1% TritonX-100, 10% glycerol) and with LiCl buffer (25mM Tris-HCl, pH 7.8, 250mM LiCl, 0.5% NP-40, 0.5% Na-deoxycholate, 1mM EDTA, 10% glycerol). The immunoprecipitated nucleic acid was eluted from the beads at 37°C in 100mM Tris-HCl, pH 7.8, 10mM EDTA, 1% SDS for 15 min. The eluted nucleic acid was purified using the DNeasy Kit (Qiagen) and analyzed by qPCR with strong-stop primers (primers P_R_ and P_U5_) or, analyzed by RT-qPCR with the P_R_ primer and another primer that anneals to nucleotides 39 to 57 in tRNAPro (P_tRNA_) (13). For the nuclease treatments, the nucleic acids were treated at 37°C with 50 U RNase A (Thermo) for 20 min in the presence of 300 mM NaCl, 4 U DNase I (Roche) with the reaction buffer provided with the enzyme for 20 min, or 3 U of RNase H (Thermo) for 20 min in the reaction buffer provided with the enzyme. Samples were purified using the DNeasy Kit (Qiagen) and the nucleic acids were subjected to qPCR or RT-qPCR analysis as described above.

### Proximity Ligation Assays

NIH3T3 or BMDC cells were seeded on glass coverslips in 24-well plates, infected with MLV (MOI=1) for 4 hr, fixed with 4% paraformaldehyde–PBS, and permeabilized with 0.25% Triton X-100–PBS. Blocking and staining were performed with Duolink in situ PLA probes and detection reagents (Sigma-Aldrich). Images were taken by using the Z-stacking function at 40X on a Keyence BZ-X710 microscope and analyzed with the BZ-X analyzer. PLA dots were counted manually in a blind fashion. Images were scored for spot localization in the nucleus (pink spots in the nucleus), nuclear membrane (red spots at the nucleus edge), and cytoplasm. Positive cells with spots in nucleus, nuclear membrane, or cytoplasm were normalized to the number of cells in the pictures. The primary antibodies used were anti-DNA-RNA Hybrid, clone S9.6 (Sigma-Aldrich), Polyclonal goat anti-MLV P10 antibody (NCI Repository), Polyclonal goat anti-MLV RT antibody (NCI Repository), anti-MLV-p30 (rat monoclonal, Hybridoma R187 ATCC®-CRL1912™), anti-NUP358 (sc-74518, Santa Cruz), and anti-NUP62 (610498, BD Biosciences).

### RNAi knockdown

For the depletion of nucleoporins in NIH3T3s and BMDCs, siRNAs from Ambion were used (catalog no: 4390771; NUP358: s72722; Nup62: n424643; Nup88: s72105; Nup153: s104224; Nup214: s105773). Briefly, BMDCs were transfected using the reverse-transfection method of the Lipofectamine RNAi MAX reagent (Invitrogen). siRNA depletion was carried out for 48 h prior to infection with MLV. RNA was isolated using the RNeasy minikit (Qiagen). RT-qPCR was performed using the GoTaq 1-step RT-qPCR system (Promega). Knockdowns were verified using primers described in Table I, for each nucleoporin, and protein expression was measured by immunofluorescence microscopy.

**TABLE I:**
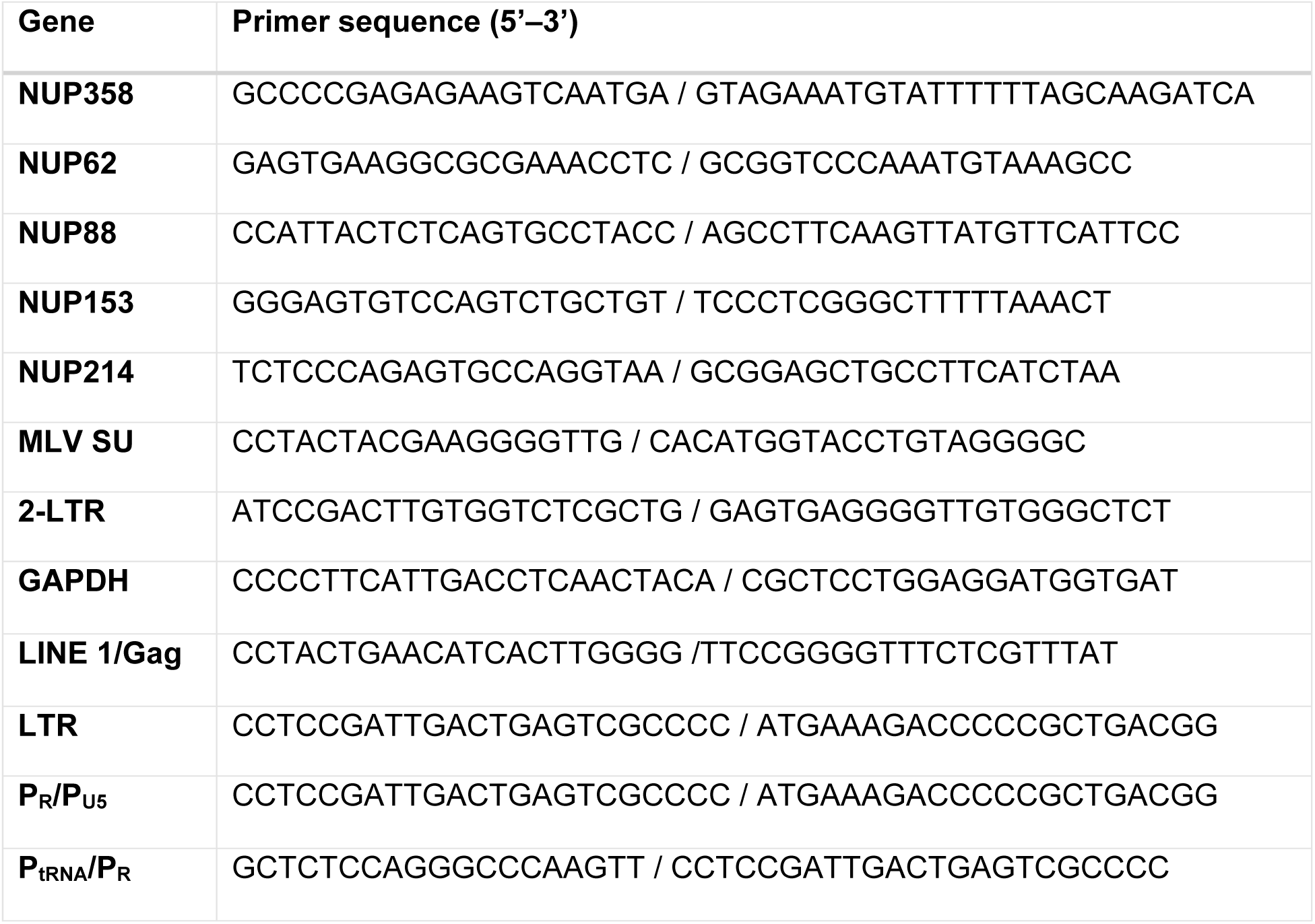
RT-qPCR primers used to evaluate cell and viral nucleic acid levels.

### Immunofluorescence microscopy

Primary antibodies used were mouse anti-NUP358 (sc-74518, Santa Cruz), mouse anti-NUP62 (610498, BD Biosciences), rabbit anti-NUP88 (AB231187, Abcam), rabbit anti-NUP153 (E316Z, Cell signaling), and rabbit anti-NUP214 (AB70497, Abcam). Goat anti-mouse IgG Alexa Fluor 488-conjugated (Invitrogen, A-11001) and goat anti-Rabbit IgG Alexa Flour 568-conjugated (Invitrogen, A-11011) secondary antibodies were used. Images were acquired using a Keyence BZ-X710 at 40X magnification. Data analysis was performed with BZ-X Analyzer software. The nucleoporin fluorescence analysis was calculated by averaging the total fluorescence intensity per total number of cells.

### Statistical analysis and data deposition

Data represent the averages of at least 3 independent experiments or as otherwise indicated in the Fig. legends. Statistical analysis was performed using GraphPad Prism 10.0.0 software. Tests used to determine significance are indicated in the figure legends. Raw data for all figures are deposited as a Mendeley dataset at xxxxxx.

## Acknowledgements

We thank David Ryan for help with mouse breeding and Monica Roth for the MLV p12 mutant plasmid.

## Author Contributions

Funding acquisition: Susan Ross

Investigation: Karen Salas-Briceno, Wenming Zhang, Susan Ross

Methodology: Karen Salas-Briceno, Wenming Zhang, Susan Ross

Project administration: Susan Ross

Supervision: Susan Ross

Writing – original draft: Karen Salas-Briceno, Wenming Zhang, Susan Ross

Writing – review & editing: Karen Salas-Briceno, Wenming Zhang, Susan Ross

---

**S1 Fig.**
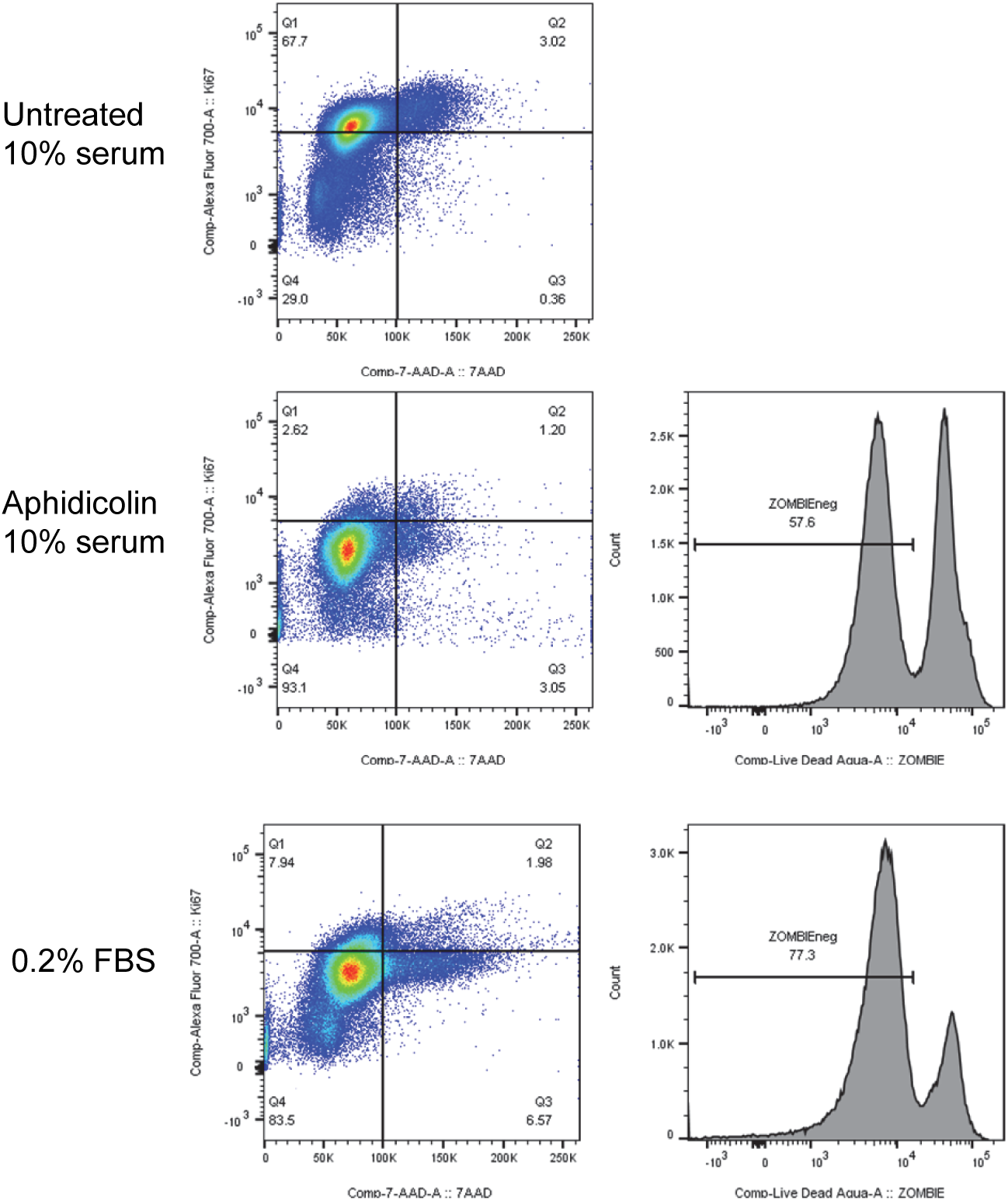
Aphidicolin-treated NIH 3T3 cells. Cell cycle analysis after aphidicolin or low serum treatment. To the right is shown the percentage of ZOMBIE-stained cells after each treatment.

**S2 Fig.**
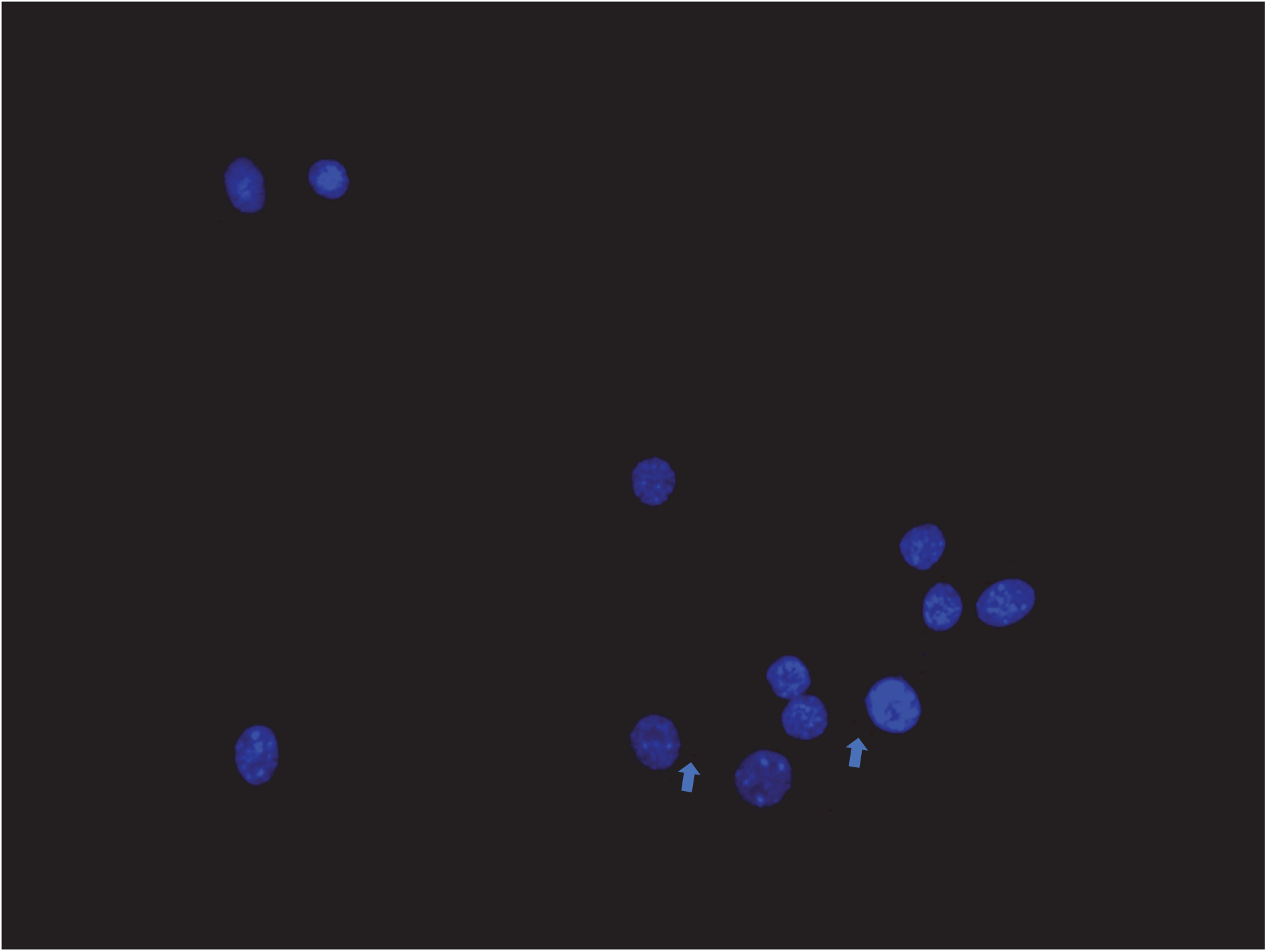
PLA showing RT-S9 interaction in uninfected NIH3T3 cells.

**S3 Fig.**
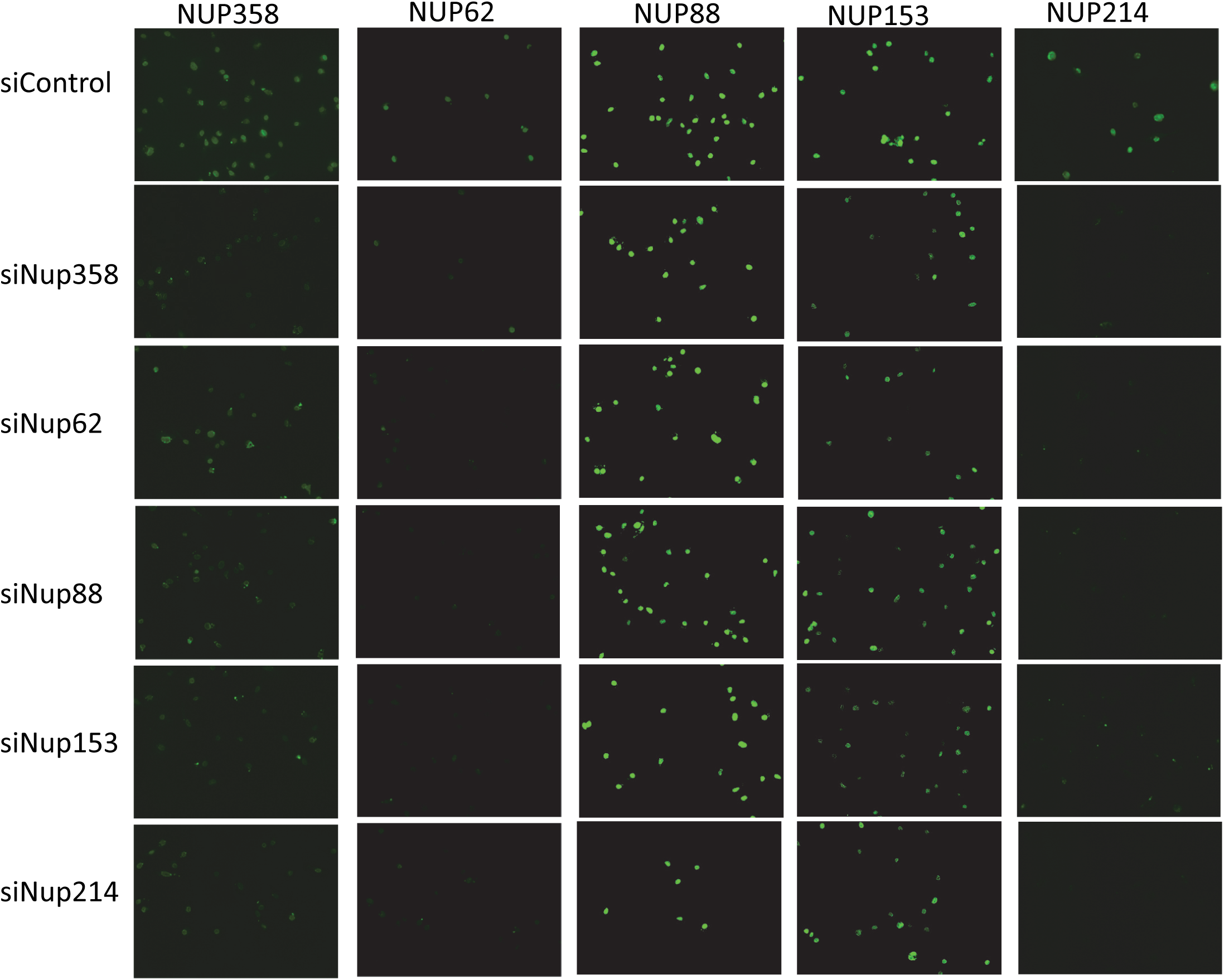
Knockdown of one NUP decreases expression of the other NUPs in BMDCs. BMDCs were transfected with the indicated siRNAs and immunofluorescence with antibodies to the different NUPs was performed. Representative images are shown.

**S4 Fig.**
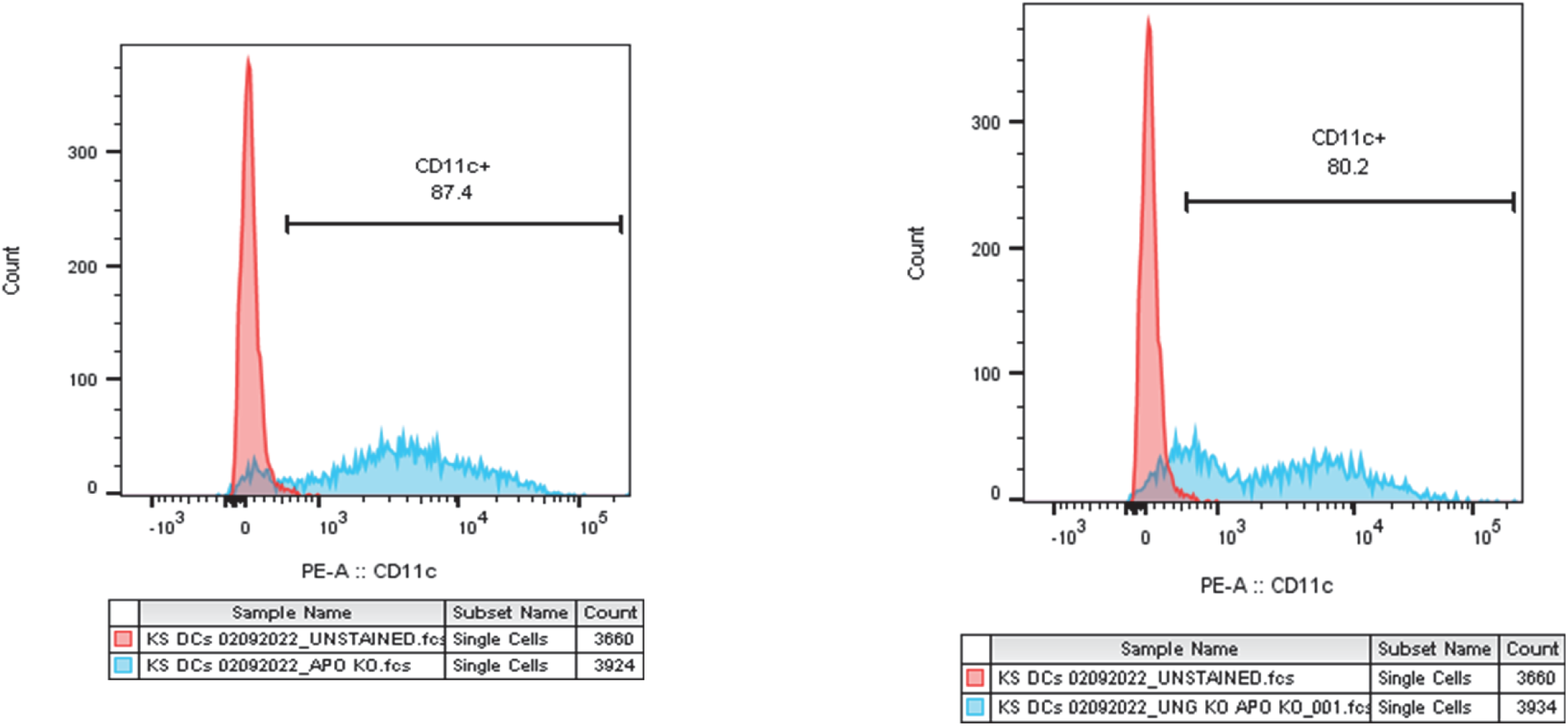
Purity of dendritic cells. Representative FACS histograms for different isolations are shown.

**S5 Fig.**
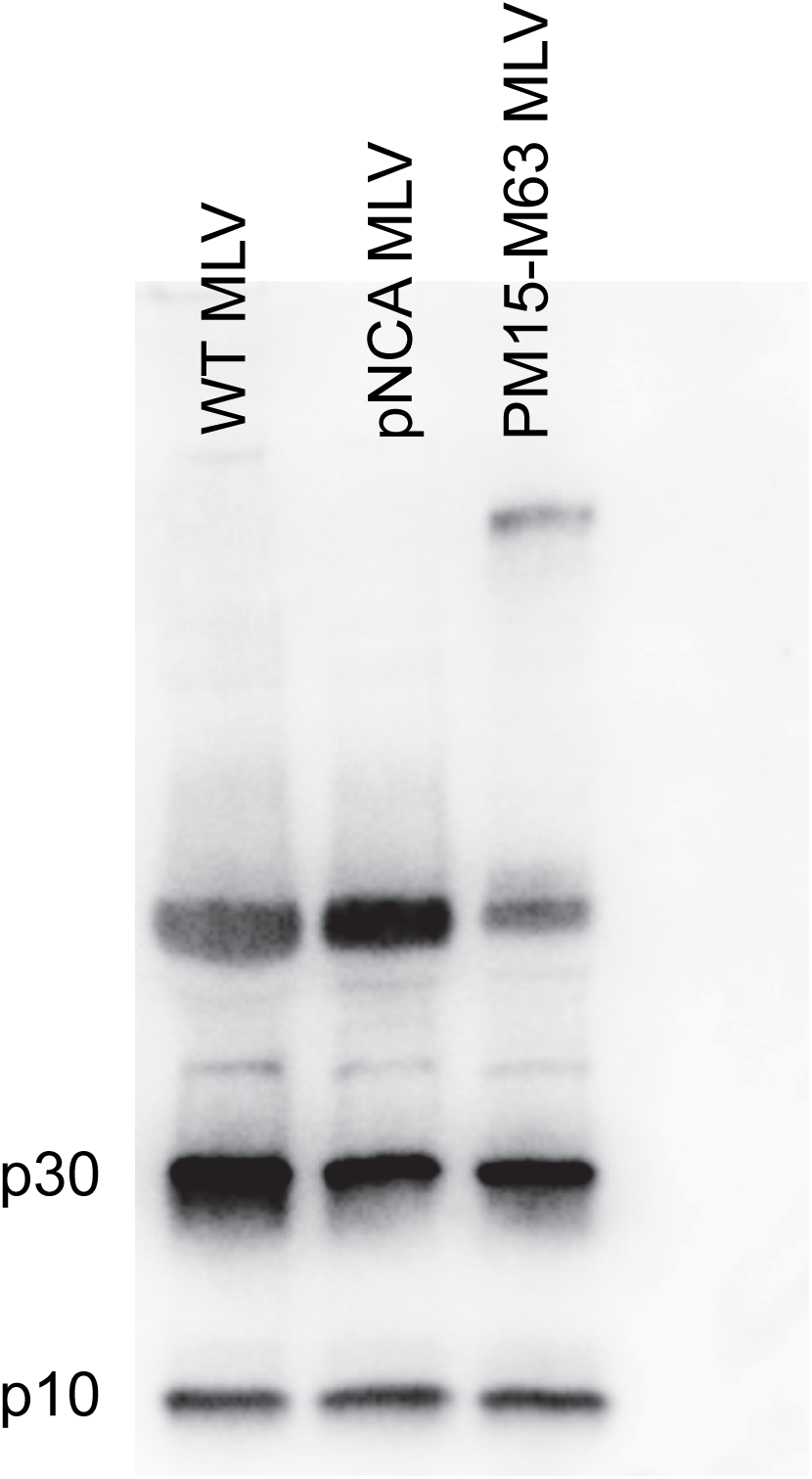
Western blots of recombinant viruses. Equal amounts of virus, were subjected to western blot analysis with anti-MLV antisera.

